# Genotyping of Inversions and Tandem Duplications

**DOI:** 10.1101/056432

**Authors:** Jana Ebler, Alexander Schönhuth, Tobias Marschall

## Abstract

**Motivation:** Next Generation Sequencing (NGS) has enabled studying structural genomic variants (SVs) such as duplications and inversions in large cohorts. SVs have been shown to play important roles in multiple diseases, including cancer. As costs for NGS continue to decline and variant databases become ever more complete, the relevance of genotyping also SVs from NGS data increases steadily, which is in stark contrast to the lack of tools to do so.

**Results:** We introduce a novel statistical approach, called DIGTYPER (Duplication and Inversion GenoTYPER), which computes genotype likelihoods for a given inversion or duplication and reports the maximum likelihood genotype. In contrast to purely coverage-based approaches, DIGTYPER uses breakpoint-spanning read pairs as well as split alignments for genotyping, enabling typing also of small events. We tested our approach on simulated and on real data and compared the genotype predictions to those made by *DELLY*, which discovers SVs and computes genotypes. DIGTYPER compares favorable especially for duplications (of all lengths) and for shorter inversions (up to 300 bp). In contrast to DELLY, our approach can genotype SVs from data bases without having to rediscover them.

**Availability:** https://bitbucket.org/jana_ebler/digtyper.git

## 1 Introduction

As of today, several population-scale sequencing projects have been finalized (The 1000 Genomes Project Consortium, 2015; The Genome of the Netherlands Consortium, 2014; The UK10K Consortium, 2015). These projects have revealed an overwhelming amount of new genetic variants and provide the basis to gain deeper insight into the principles of evolution and the association of variants with phenotypes, where disease risks play a particularly important role.

A crucial step in integrating variants into studies on evolution and disease is to genotype and phase them. That is, one has to first determine their zygosity status (genotyping) and then partition the alleles at heterozygous loci into two groups reflecting the two parents of the individual (phasing). Thereby, the accuracy of the second step crucially hinges on the first step. This implies that one has to operate at utmost accuracy when genotyping—only if genotypes have been determined carefully, variants can finally serve the purposes of downstream studies.

Genotyping variants from next-generation-sequencing (NGS) data, however, can pose involved computational challenges. While geno-typing single nucleotide polymorphisms (SNPs) from NGS data already is a routine procedure, genotyping more complex and larger variants is not. Recent advances have pointed out how to do this for shorter (≤ 20–30 bp) insertions and deletions (indels), larger deletions and mobile element insertions (The 1000 Genomes Project Consortium, 2015; Hehir-Kwa *et al.*, 2016). However, the majority of inversions, duplications and translocations are still lacking sound models that allow to determine their genotypes from NGS data. So far, only a few such variants have made their way into phased reference panels (The 1000 Genomes Project Consortium, 2015; Hehir-Kwa *et al.*, 2016). The distinguishing feature of the already phased such variants usually is that they stem from genomic regions that exhibit particularly favorable sequence context. This, however, applies for only little of them. The major part of inversions, duplications and translocations stem from genomic regions that are “inaccessible”, difficult to analyze by short read data. Therefore, simple ad-hoc approaches to genotyping such variants do not work.

In this paper, we will provide statistical models and efficient computation schemes that allow to genotype tandem duplications and inversions even though the corresponding read data does not necessarily stem from highly “accessible” genomic regions. The challenge in this is to control the statistical uncertainties that affect the NGS data that give evidence of such variants. In the first place, aligning the affected reads poses particular difficulties for short read alignment programs such that many of their alignments remain ambiguous. As a result, many likely variant affected reads are uncertain in terms of their placement. If an alignment is incorrect, that is, the read does not even stem from the variant region in question, it provides no insight into the zygosity status whatsoever. Second, even if correct, one can often interpret an alignment in multiple ways, which may lead to contradicting statements about the existence of variants.. The latter case is due to the fact that fragment length can vary and/or the ambiguous placement of alignment breakpoints, although the alignment overall indicates the correct placement, among other issues.

Here, we have been inspired by the models presented by Hehir-Kwa *et al.* (2016, Supplement, Section 5.2), which have led to genotypes of high accuracy for deletions and insertions. These models follow the principle to infer the genotype that is most likely in terms of the read data that supports it. A major problem of such a maximum likelihood (ML) approach for computing genotypes from NGS read data is that a naive evaluation of all relevant reads, together with their uncertainties, results in exponential runtime algorithms. The exponential “explosion” in runtime is a common problem when taking uncertainties into additional account. Here, for the first time, we present a computation scheme that has runtime linear in the number of the relevant data, even if affected by uncertainties.

An additional advantage of the rigorous statistical approach presented is that the genotype likelihoods computed, that is the probabilities that a variant is absent, heterozygous or homozygous, are highly reliable. Unlike for simple ad-hoc counting strategies, our approach does not get confused by the uncertainties involved. As usual, the genotype likelihoods can be further used for filtering, thereby controlling the quality level one intends to operate on in downstream analyses, which, in particular, includes computational phasing pipelines. Very often, the underlying data may not allow to correctly distinguish between two genotypes, because the uncertainties affecting the data are too large, or there is too little coverage. In this case, the genotype likelihoods should reflect such situations, such that downstream method can come to the appropriate conclusions.

**Related Work.** In comparison to discovery, there are relatively few approaches that allow to genotype non-SNP genetic variants. Most of these approaches specialize in genotyping insertions and/or deletions of varying length. We cite a few prominent and widely used such approaches and refer the interested reader to Lin *et al.* (2015) for further references. Platypus (Rimmer *et al.*, 2014) generally focuses on smaller indels, but, by making use of local assembly, can also genotype larger ones. Pindel (Ye *et al.*, 2009) also offers a basic procedure for genotyping indels of length up to 50-60 bp. GenomeStrip (Handsaker *et al.*, 2011) has been in use at the 1000 Genomes project (The 1000 Genomes Project Consortium, 2015) for genotyping large deletions. MATE-CLEVER (Marschall *et al.*, 2013) can genotype midsize and long deletions and has been in use at the Genome of the Netherlands project (Hehir-Kwa *et al.*, 2016); the most recent version implements the framework that inspired the present paper.

There are also well-known methods that at least in principle allow to genotype inversions. First, Cortex (Iqbal *et al.*, 2012) generally offers geno-typing for various kinds of non-copy-number variants using a colored de Bruijn graph approach, but has not been evaluated for inversions. GASV-PRO (Sindi *et al.*, 2012) offers to discover inversions and a sound statistical model for genotyping variants in general, however, the genotyping option has not been evaluated for inversions.

The only method we are aware of (and believe is the only one) that offers to genotype tandem duplications and inversions at sufficiently high quality, as per publicly available evaluation experiments, since being in use at the 1000 Genomes project, is DELLY (Rausch *et al.*, 2012). This explains why we will focus on DELLY as a comparison partner in our benchmarking experiments.

**Contributions.** In this paper, we describe a novel statistical framework called *DIGTYPER* (*Duplication and Inversion GenoTYPER*) that allows to genotype inversions and tandem duplications while keeping control of the uncertainties that affect the corresponding read data. We provide an efficient algorithm, by which to compute the genotype likelihoods while taking uncertainties into account. The algorithm has runtime linear in the number of the affected read alignments, which overcomes a classical obstacle when dealing with uncertain data, because naive approaches lead to exponential runtime. As results, we demonstrate that we achieve very favorable results in comparison to DELLY (Rausch *et al.*, 2012), which, to our knowledge, is the only approach available that allows to genotype tandem duplications and inversions at operable accuracy levels. On simulated data, we clearly outperform DELLY for tandem duplications. For inversions, we are on a par with DELLY for longer inversions. DELLY, however, cannot genotype shorter inversions, for which we achieve performance rates similar to those for longer inversions. Last but not least, unlike DELLY, our method is able to acknowledge too large uncertainties, in which case it will correctly point out the corresponding ambiguities.

## 2 Methods

Our goal is to compute genotypes for given inversions and tandem duplications based on aligned sequencing reads. To this end, we adapt a procedure that we employed previously to genotype insertion and deletions, see Hehir-Kwa *et al.* (2016, Supplement, Section 5.2). For every variant to be genotyped, we consider all reads from the corresponding region. This set of reads, referred to as 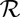, is used to determine the genotype for the respective variant. The goal is to compute probabilities for the three possible genotypes *G_i_*, for *i* = 0, 1, 2, given all reads 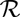, where *G*_0_, *G*_1_, *G*_2_ represent that the variant in question is absent, heterozygous or homozygous, respectively. We use the reads from 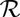 to compute a posterior probability for each genotype. Finally the genotype for which the probability is highest is the result.

For each read 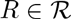, let ℙ(*A*^+^(*R*)) denote the probability that its alignment is correct and let ℙ(*A*^−^(*R*)) denote the probability that it is not. Then, ℙ(*G_i_*|*A*^+^(*R*)) is the probability for genotype *G_i_* under the assumption that the read is mapped correctly and ℙ(*G_i_*|*A*^−^(*R*)), the probability for a genotype under the assumption that the alignment of the read is wrong. Using Bayes’ theorem and the assumption of constant priors over the genotypes, the probability for a particular genotype given all reads can be expressed as follows (Hehir-Kwa *et al.*, 2016):

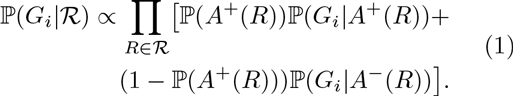

Note that this expression can be evaluated in time linear in 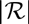, and hence avoids to explore all 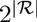 possible combinations of correctly/wrongly mapped reads, which would result in exponential runtime. In the following, we derive the terms needed to evaluate (1). We set

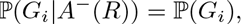

where ℙ(*G_i_*) expresses our prior belief in genotype *G_i_*, because if the considered read does not stem from the region it does not give any information about the genotype.

The probabilities for the alignment of the read to be correct, ℙ(*A*^+^(*R*)), and to be incorrect, ℙ(*A*^−^(*R*)), can be obtained as follows.

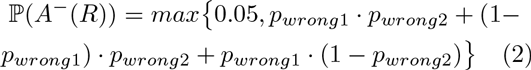

where

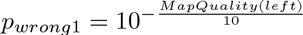

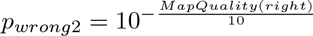

are the probabilities for the two read ends of a read pair not to be mapped correctly, so *p_wrongi_* gives ℙ(*A*^−^(*R*)) for the left end and analogously *p_wrong2_* for the right end of the considered read pair. The *MapQuality* of a read is directly taken from the input BAM file. By considering the maximum in Equation 2, we ensure to never fully “trust” a read alignment, and hence account for mapping uncertainties not recognized by the read aligner. Finally ℙ(*A*^+^(*R*)) is computed as 1 – ℙ(*A*^−^(*R*)).

The only yet unspecified probability to evaluate (1) is ℙ(*G_i_*|*A*^+^(*R*)). Its choice depends on whether inversions or duplications are considered, as we see in the following sections.

### 2.1 Approach for Inversions

For inversions two cases must be distinguished, depending on the positions of the mapped reads in the dataset. In both cases, reads supporting the variant and such that support the reference are used for the computation of the likeliest genotype.

#### 2.1.1 Reversed Read Evidence

In this case one end of the paired-end read is mapped left or right of the inversion and the other end is mapped completely inside the inversion. Under the assumption of no alignment uncertainty, a read stems from a sequence that contains the inversion if and only if the orientation of the read end located inside the inversion is the same as for the other read end. This means the read end in the inversion was reversed when being mapped to the reference. See Figure 1 for an example. Such a read supports the presence of the considered inversion in the sequence. Let *R*_1_ be a read that supports the inversion. According to Bayes’ theorem and the assumption of constant priors, it holds that ℙ(*G_i_*|*A*^+^(*R*_1_)) α ℙ(*A*^+^(*R_1_*)|*G_i_*). The latter term can be computed as follows. ℙ(*A*^+^(*R_1_*)|*G_0_*) = 0, because if the inversion is absent the read cannot stem from the region. Then 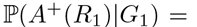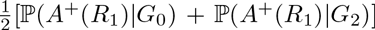, reflecting the case that one randomly picks one of the two chromosomal copies with only one containing the inversion and then generates the read from it. These considerations lead to the expression

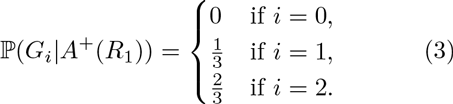

**Figure 1:**
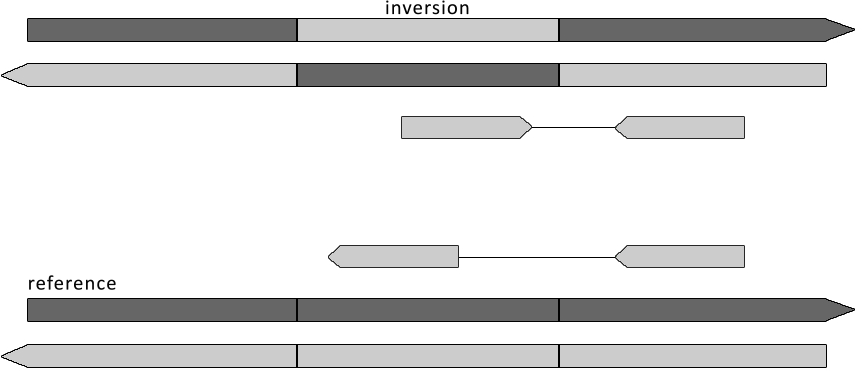
Reversed Read Evidence. Above the original read is shown and below the read is mapped to the reference. Note that the end that stems from in between the inversion breakpoints changes its orientation when being mapped to the reference.

If the orientation of the read mapped in between the inversion breakpoints is not changed, the read must stem from a sequence without the inversion. Let *R*_2_ be such a read. This case is treated analogously to the previous one, but this time we have ℙ(*A*^+^(*R_2_*)|*G_2_*) = 0, leading to

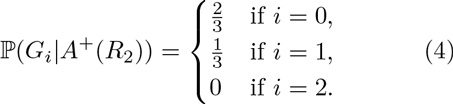

#### 2.1.2 Split Read Evidence

Read pairs for which one read end stretches across one of the inversion breakpoints cannot be mapped by standard read mappers. To leverage these reads for genotyping, we extended the read mapper LASER (Marschall,T. and Schönhuth,A., 2013) to detect inversion-type split alignments (see Figure 2). Since LASER is based on partial banded alignments that extend seed hits, implementing this feature only required to combine anchor alignments of opposite directionality (showcasing the power of this technique). When using option --**inversions**, LASER output these split reads encoded as IV tags in the BAM output.

**Figure 2:**
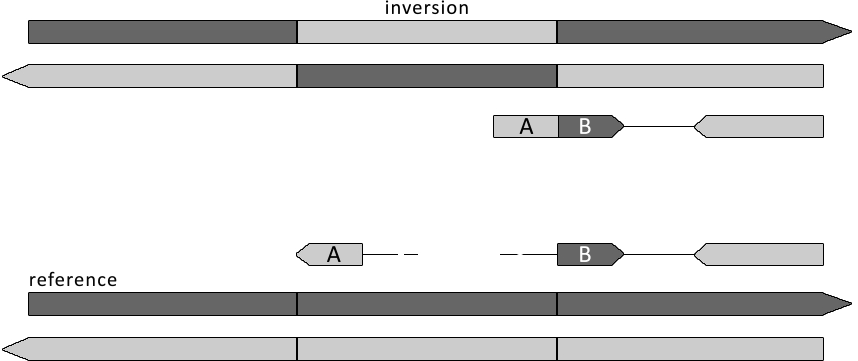
Split Read Evidence. The light part of the left read end is reversed after mapping while the rest is not changed. There will be a long gap between the light and the dark part of the left read end.

For genotyping, we evaluate these tags and decide whether a (split) alignment across an inversion breakpoint supports the inversion or the reference. After being mapped to the reference sequence, the part of this read end located inside the inversion breakpoints (:= *A*) is reversed, while the other part outside the breakpoints (:= *B*) is not, as illustrated in Figure 2. The distance between the alignments of *A* and *B* equals *length*(*inversion*) — *length*(*A*). To support the inversion the following requirements have to be fulfilled besides the reversed orientation of *A*: One end of part *B* must agree with the inversion breakpoint the read stretches over and one end of part *A* has to agree with the other inversion breakpoint. The read supports the inversion if and only if these requirements are fulfilled. Otherwise it supports the reference sequence. Just as in Section 2.1.1, the probability ℙ(*G_i_*|*A*^+^(*R*_1_)) for the genotype *G_i_* is computed using Equation (3) and Equation (4) for reads supporting inversion allele and reference allele, respectively.

For split-reads ℙ(*A*^−^(*R*)) is computed as

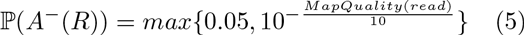

Here *read* describes the read end which stretches over the breakpoint. Since for split reads only these read ends are considered, only the probability that such an end is wrong, is taken into account. Again, ℙ(*A*^+^(*R*)) is computed as 1 – ℙ(*A*^−^(*R*)).

### 2.2 Approach for Duplications

To genotype duplications, we use a statistical framework that considers all read pairs with at least one read end aligned to lie completely inside the duplication. Figure 5 shows how such read pairs can be placed on the originating duplication allele and to how the resulting alignments look like. Compared to inversions, we now have to overcome an additional complication: No read pair gives direct evidence of the reference allele. All read pairs that originated from the reference allele could potentially also have originated from the duplication allele. Read types B and D in Figure 5 are examples of this. For inversions, that was not the case and we could restricted our attention to read pairs that can uniquely be determined to either stem from the variant allele or from the reference allele.

**Figure 3:**
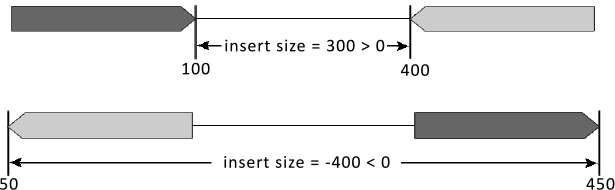
Definition of the insert size of a paired-end read.

In the following, reads that unambigously support a duplication are denoted as *supporting* and read pairs that can have originated from both alleles (reference/duplication) are denoted as *neutral*. In order to genotype duplications, the main idea is to consider the proportion of supporting and neutral reads, which can be achieved within the same framework as for inversions. Again, our goal is to compute ℙ(*G_i_*|*A*^+^(*R*)) in order to evaluate Equation (1).

We will approach short and long duplications separately since they are qualitatively different in terms of read types shown in Figure 5. For short duplications, read pairs of types A, B, C, D, and E exist. Read pairs of type F do not exist when the duplication is smaller than the fragment size (i.e. insert size plus read lengths). For long deletions, read pairs of type F are present but types A and C do not exist. Note that read pairs of type E exist for short and for long duplications, but their relative placement on the reference genome depends on the duplication length. For a deletion longer than the fragment size, the order of forward and backward read ends on the reference gets reversed as can be seen in 5. We give a precise definition of “short” and “long” after introducing some notation in the following section.

#### 2.2.1 Insert Size Evidence

For duplications, the distance of the alignments of the two read ends in a read pair can be leveraged for genotyping. Consider a scenario where a read pair stems from the duplication allele and the left read end lies outside and the right read end lies inside the duplication. The right read end could have originated from the first copy or the second copy of the duplication in the donor genome. In the latter case, we observe a reduced distance of the aligned read pair, as illustrated in Figure 5, read type A. In slight abuse of terminology, we refer to the distance of the two aligned reads as *insert size*, depicted in Figure 3. The insert size is computed by subtracting the end position of the read end which is mapped to the forward strand from the start position of the read end which maps to the reverse strand. In case the orientations of the read ends are reversed, this value is negative (Figure 3).

We adopt the common assumption (Marschall *et al.*, 2012) that the insert size follows a normal distribution under the null hypothesis (i.e. when the read pair stems from the reference allele and has been mapped correctly). Mean and standard deviation of this distribution can be robustly estimated from the aligned reads (Marschall *et al.*, 2012). We hence assume these quantities to be known and denote them as *µ* and *σ* in the following. A normal distribution with this mean and standard deviation is written as 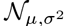.

Let *ℓ* denote the length of the duplication to be genotyped, which we define as the length of the repeat unit that is duplicated and therefore occurs twice in the genome. Read pairs spanning a copy of the duplicated sequence, such as read pairs A and C in Figure 5, will have an observed insert size distributed according to 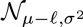, whereas read pairs from the reference allele exhibit insert sizes distributed according to 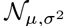.

#### 2.2.2 Short Duplications

We call a duplication *short* when its length *ℓ* is smaller than *µ* + *len*(*ReadEnd*). This implies that read pairs of type F (Figure 5) are very unlikely to exist.

The most important evidence for short duplications comes from read types A, B, C, and D. They are characterized by one read end being aligned completely inside the duplication and the other read end being aligned (at least partially) outside the duplication. We call the read completely inside the duplication *anchor* read. As outlined above, these read pairs can either lead to an observed insert size distribution of 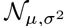 (for types B and D) or of 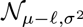 (for types A and C). To derive ℙ(*G_i_*|*A*^+^(*R*_1_)), we consider how these read pairs can be generated, as illustrated in Figure 4. The probability of whether a given read pair has originated from the reference or from the duplication allele obviously depends on the genotype. While they equal 1 for homozygous genotypes, a heterozygous genotype leads to probabilties to stem from the reference allele or duplication allele of 1/3 and 2/3, respectively. The duplication allele is twice as likely because the anchor read is completely inside the duplicated region which exists twice on the duplication allele. In case a read came from the reference allele, we observe an insert size distribution of 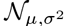. In case it came from the duplication allele, two scenarios are possible: either the anchor read originated from its first copy or from it originated from its second copy, leading to either an observed insert size distribution of 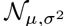 or of 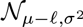. The whole process is illustrated in Figure 4.

**Figure 4:**
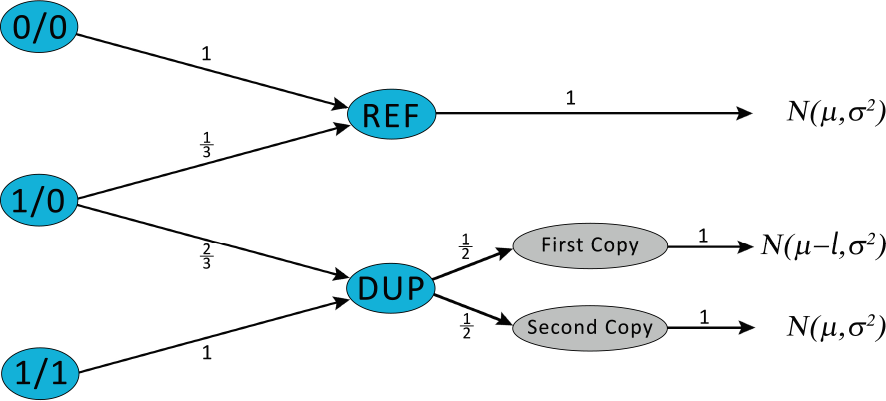
Link between genotypes (left column) and distributions of an observed insert size (right column) mediated by the allele the read pair stems from (middle, blue) and, in case of the duplication allele, by the copy of the duplication the anchor read originates from (middle, grey). We show the scenario where the left read end is the anchor; if the right read end is the anchor, the roles of “first copy” and “second copy” are swapped. Edge labels indicate probabilities.

Each path from left to right in Figure 4 contributes to the probability that a given genotype in the left column gives rise to read pairs with the observed insert size distribution given in the right column. By summing up all paths for each geno-type, we obtain

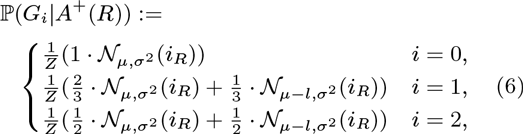

where *i_R_* denotes the insert size observed for read *R* and *Z* is a normalization factor given by

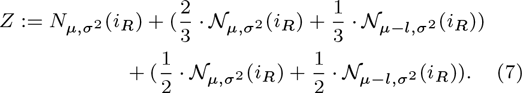

Read pairs of type E are processed using the same formula. Summing over all read pairs by plugging Equation (6) into Equation (1) yields the sought genotype likelihoods.

#### 2.2.3 Long Duplications

We call a duplication *long* when its length *ℓ* is longer than or equal to *µ* + *len*(*ReadEnd*). This makes it possible that read pairs of type F exist, which require some extra attention. One observes twice the number of such read pairs per duplication allele than per reference allele and their expected number grows linearly with the duplication length (in fact constituting the signal that coverage-based copy number estimation tools use). Like for short duplications, we seek to only use those read pairs that span a duplication breakpoint, either start or end of the duplicated region or the internal breakpoint between the two copies of the duplication. That is, we want to use read pairs of type E but exclude those of type F. For long duplications, however, this distinction can be made with very good accuracy based on the observed insert size (for values of *µ*, *σ*, and *len*(*ReadEnd*) common in practice). Read pairs of type E have reversed orientations or at least they overlap after they have been mapped to the reference. After discarding read pairs of type F, the genotyping proceeds in the same way as for short duplications. The sum over all reads 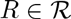 in Equation (1) runs over all read pairs that have an anchor read mapped completely inside the duplication. It is important to note that read pairs of type E have *two* anchor reads and hence need to be counted twice in this sum. This ensures that the expected number of counted read pairs of type E equals the expected number of read pairs of types B and D. Note that observing reads of types A or C becomes increasingly unlikely for growing duplication lengths.

### 2.3 From Likelihoods to Genotypes

After genotype likelihoods have been computed as explained above, the likeliest genotype is reported as result. DIGTYPER also outputs genotype likelihoods as phred-scaled posterior probabilities when writing a VCF file. That is, —10 · log_10_(*p_i_*) is reported as genotype likelihood for genotype *i* with posterior probability *p_i_*. The difference of the phred-scaled posterior of the likeliest and second-likeliest genotype is used to decide whether to report a genotype at all or *./.* to indicate too large ambiguity; the default threshold for this difference is set to 20 and used throughout the evaluation. When this filter has not been passed but the phred-scaled posterior for genotype *0/0* (homozygous in the reference allele) is below a user-specifyable threshold (default set to 20), then genotype *1/.* is reported to indicate that at least one alternative allele is present, i.e. the genotype is believed to be either *1/0* or *1/1* but the data is insufficient to distinguish these two cases.

## 3 Results

We evaluated our algorithm on simulated data and on a data set provided by the *Genome in a Bottle Consortium* (*GIAB*) (Zook *et al.*, 2015). We compare our algorithm with DELLY (Rausch *et al.*, 2012), which, as pointed out above, is the only approach that has been publicly evaluated on inversions and tandem duplications. DELLY cannot be run in “genotyping-only mode”, but can only genotype its own discoveries, which explains why we can only evaluate DELLY on its own calls in the following. For head-to-head comparison with DELLY, we evaluate our own method on only DELLY calls. As one can expect to see variant databases being steadily filled with inversions and tandem duplications in the short and midterm future, there is an obvious need for tools that do not depend on their own discovery functionalities. So, we will further also evaluate our own method on all variants known relative to the respective evaluation scenario so as to gauge the extent of variants that can be geno-typed by a discovery independent approach.

**Figure 5:**
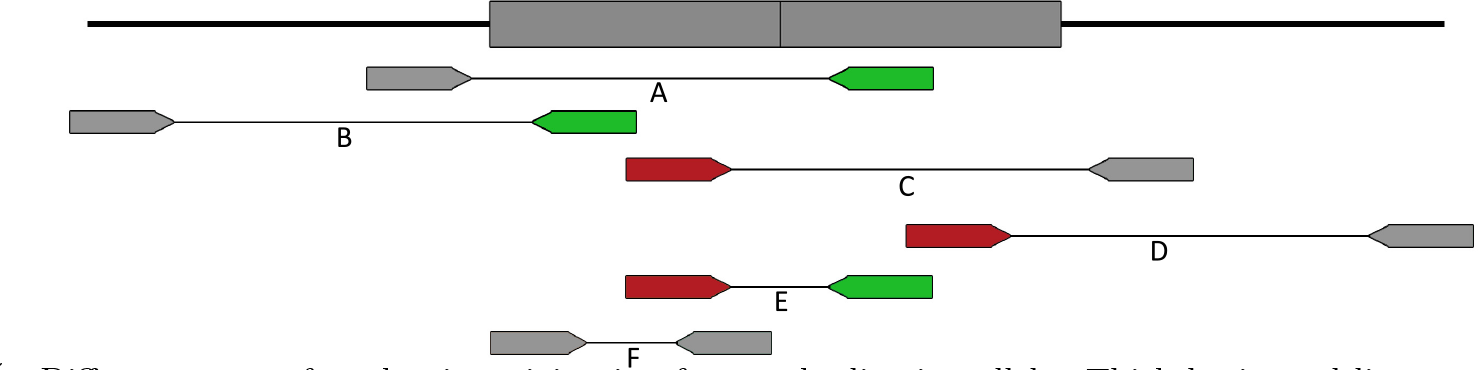
Different types of read pairs originating from a duplication allele. Thick horizontal line on top indicates the donor genom with the duplication shown as grey boxes. Forward anchor reads are shown in red and reverse anchor reads in green. Mapping supporting read pairs A, C, and E to the reference gives rise to shorter observed insert size, while the insert size of neutral read pairs B and D agrees with the null distribution 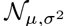.

### 3.1 Simulated Data

For generating simulated data, we used the reference sequence of human chromosome 1 (version: hs37d5). We inserted inversions and duplications of varying lengths into this chromosome, with neighboring inserted variants seperated by one million bp of reference sequence. We then used SimSeq (Earl *et al.*, 2011), a well known read simulator used in the Assemblathon (citation) for generating read data from the resulting sequence. The mean *µ* of the length of the generated fragments was 550 bp, at a standard deviation *σ* of 140. Read ends were of length 148 bp. This mimicks the parameters from the real GIAB dataset, so as to have a realistic simulation scenario. We used *bwa* (Li,H. and Durbin,R., 2009) to map the reads to the reference sequence and to create BAM files as input for the programs. Thereby, we varied the coverage and obtained datasets at coverages of 4×, 12× and 60×, all of which reflect realistic settings. While the length of the simulated inversions were 100, 300, 500, and 800 bp, the length of the duplications was set to 200, 300, 500, and 800bp—because duplications shorter than read length cannot be detected. Reads reflecting heterozygous variants were generated by simulating reads from both our simulated sequence and the reference chromosome 1, which were subsequently merged using SAM-tools (Li *et al.*, 2009).

We then ran DELLY on the generated datasets in discovery mode to generate inversion and duplication calls. The positions of these calls were then provided as input to our genotyping program. Only predictions tagged as “precise” by DELLY were considered, since for split read analysis the breakpoints should be accurate. In our evaluation experiments, we only considered DELLY variants whose center points were found to not deviate by more than 50 bp from the true center points. On these variants, we compare our genotype predictions (“DIGTYPER retype” in Figure 6) with the ones from *DELLY.* Additionally, we also evaluate our program on all variants we have inserted in our simulated data, which DELLY does not allow to do (“DIGTYPER all” in Figure 6).

#### 3.1.1 Results for Inversions

Figure 6(left) shows results for inversions. DELLY could not genotype inversions of length 100 bp, because its discovery module does not allow to detect them in the first place, which kept us from comparing our tool with DELLY on this (short) length. However, our approach delivered substantial amounts of well typed inversions if not depending on DELLY calls. This points out one first significant advantage of our tool, as it thus establishes the first approach to genotype short inversions (100-200 bp) at provably sufficiently high accuracy, in particular for coverages at least 12×. In general, our approach was superior to DELLY at low coverage, where DELLY could not genotype again due to its weakness to depend on the discovery machinery, which does not deliver sufficiently many calls at low coverage. In an overall account, our method and DELLY largely agreed on all other length ranges (starting from 500 bp) and coverage rates (starting from 12×), with DELLY yielding slighly more false predictions in comparison to our approach. Using DELLY calls, the amount of vari-correct ants for which a correct prediction could be given was between 10 and 30% for inversions of length 300 and 500 bp, while it was between 35 and 40% for the longer ones. When using the positions of all, true inversions we had inserted into chromosome 1, we found the amount of correct predictions to raise to more than 60–70% for inversions of length 300 bp and longer at coverage 4×, and to more than 80% for inversions of all length ranges at coverage rates of 12× and 60×. There is one weak spot, which is to genotype short inversions at very low coverage (4×), where only 10% could be typed correctly. However, this may point out a limitation of the data rather than a limitation of our tool. By making use of our statistical machinery, we could decide correctly in 40% of the cases that the variant existed while refusing to type, since the data uncertainties were too high to allow us to do so. while, still, we could make use of our power to rate calls too ambiguous, but still to decide whether they were present (=’1/.’) or not in 40% of the cases. In general, our program yielded very little errors, even at low coverage. In particular at low coverage, this is also due to correctly refusing to type based on the data given, and to issue genotypes ‘*1/.*’ or ‘*./.*’ in that case.

**Figure 6:**
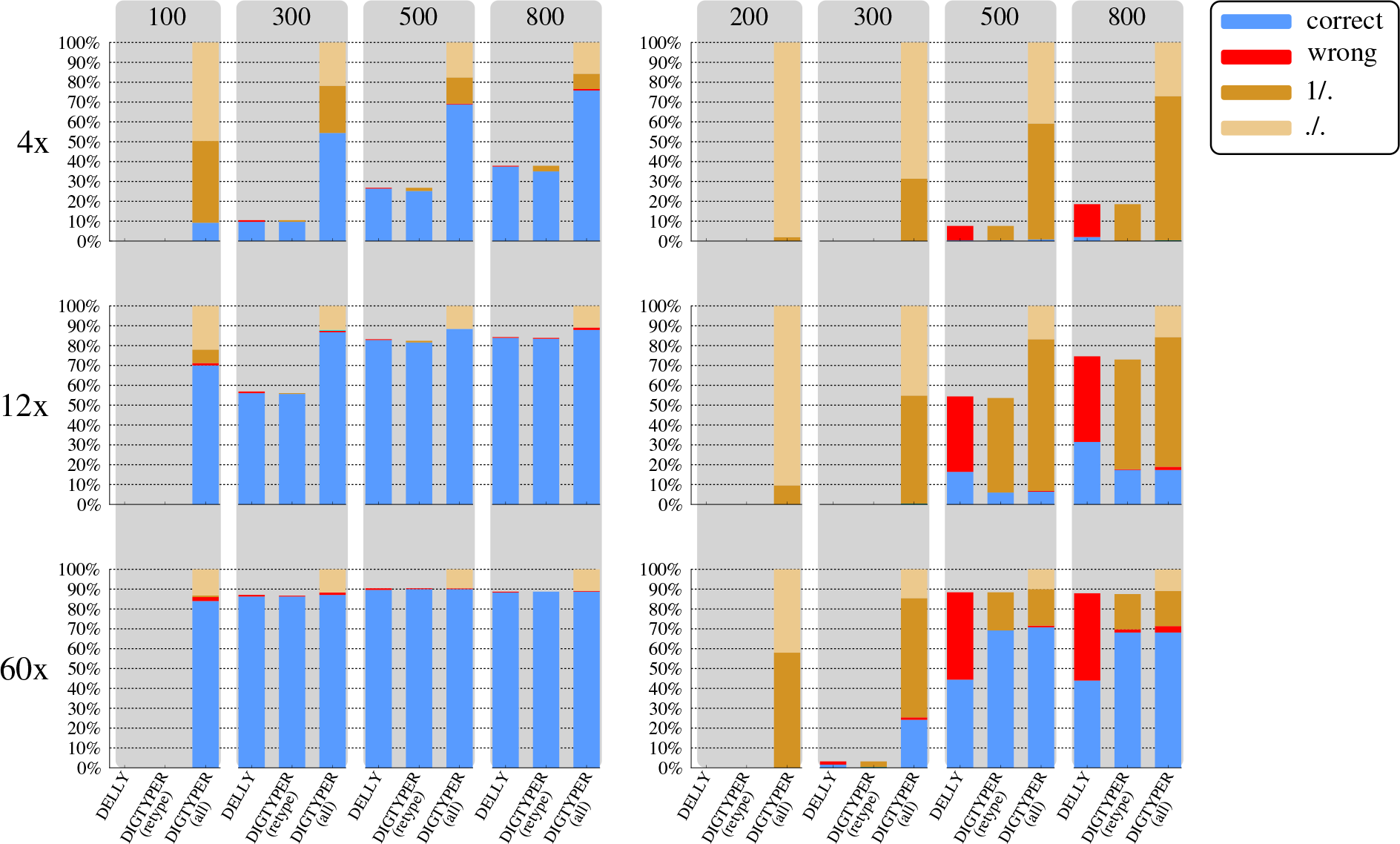
Results on simulated data for **inversions** (**left**) and **duplications** (**right**). Each triple of bars shows the amount of variants found and genotyped by DELLY, the genotype predictions made for these variants by DIGTYPER (called “DIGTYPER retype”), and the predictions made by DIGTYPER run on the full set of inserted variants without running DELLY before (“DIGTYPE all”). Results are stratified by length (grey columns, length given at the top of each column) and by coverage (rows).

#### 3.1.2 Results for Duplications

Figure 6(right) shows results for duplications. DELLY was able to discover significant fractions of implanted duplications only for lengths 500 bp and 800 bp. Comparing genotyping performance for 200 bp and 300 bp was hence not possible. These length classes are particularly challenging since only few anchor reads, which lie completely inside the duplication exist. Furthermore, such duplications are also relatively short compared to the standard deviation of the insert sizes (*σ* = 140), inducing uncertainty when distinguishing supporting from neutral read pairs. DIGTYPER recognized this uncertainty and reported *./.* and *1/.* genotypes. For longer duplications or higher coverage, the uncertainty decreased, as expected.

Even for longer duplications, DELLY was not able to distinguish homozygous and heterozygous tandem duplications, since all variants found were reported as *1/0*. This leads to a very high amount of false predictions, as evidenced by the red bars in Figure 6(right). As for inversions, the fraction of duplications discovered by DELLY was strongly influenced by the coverage. Again, by genotyping the real positions of the variants inserted there were more variants for which a genotype prediction could be given, which mimicks the application of genotyping data base variants. In contrast to the results for inversions, for duplications there was a larger amount of *1/.* predictions, especially at coverages 4× and 12×, indicating that at least one chromosomes carries the variant with a high probability. This reflects the fact that duplications are fundamentally more difficult to genotype than inversions, because no read pairs directly evidencing the reference alleles can be used. For the highest coverage of 60×, the amount of correct predictions was much higher compared to DELLY and, at the same time, also the amount of false predictions was much smaller than for DELLY.

### 3.2 *GIAB* data

We used an Illumina HiSeq dataset from *Genome in a Bottle Consortium* (Zook *et al.*, 2015) for individual HG003 of the Ashkenazi trio, which has coverage 60×. Since we lacked reliable ground truth data of genotyped inversions and duplications, we conceived the following experiment. First, we ran DELLY to discover and genotype inversions and duplications on this data set. Second, we considered all inversions and duplications reported by The 1000 Genomes Project Consortium (2015) and The Genome of the Netherlands Consortium (2014), which we refer to as *data base variants*. Note that the Ashkenazi trio is not part of either of the two projects. We genotyped these data base variants in the GIAB individual with DIGTYPER. Then, we determined for each data base variant whether it matched at least one variant discovered by DELLY (with a center point distance and length difference of up to 200 bp). Next, we determined the intersection between the sets of data base variants typed *1/1* or *1/0* or *1/.* by DIGTYPER on the one hand and the set of data base variants matching a DELLY call on the other hand. The results are shown in Figure 7.

**Figure 7:**
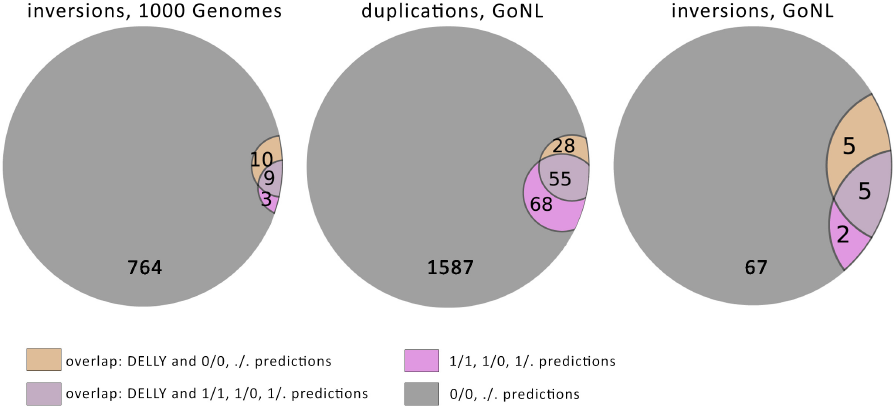
Shown are the overlaps of genotype predictions (*1/1,1/0,1/*. and *0/0, ./.* for inversions and duplications given by The 1000 Genomes Project Consortium (2015) and The Genome of the Netherlands Consortium (2014) made by our program, and the genotype predictions for corresponding variants found by DELLY. The gray areas hence represent data base variants for which DELLY and DIGTYPER agree that the variant is absent from that individual, while the dark purple areas represent variants for which both methods agree that the variant is present.

Still lacking a ground truth, we can now compare DELLY calls to DIGTYPER predictions. Since DELLY often reports multiple overlapping predictions (sometimes with different genotypes) matching the same data base variant, we did not compare genotypes, but only absence/presence signals. We want to emphasize that data base variant typed as 0/0 by DIGTYPER and not discovered by DELLY are *not* false negatives but, most likely, constitute variants simple absent in the studied individual.

From 6025 duplications given by The 1000 Genomes Project Consortium (2015), only two were deemed present (i.e. 0/1 or 1/1 or 1/.) by DIGTYPER and two were deemed present by DELLY, with one variant overlap. Because of these small numbers, we omited 1000 Genomes duplications from Figure 7, which summarizes the findings for 1000G inversions and GoNL inversions and duplications.

In all cases, most of the variants were genotyped as *0/0*. For the majority of those, no matching DELLY variant was found and therefore they are likely to be correctly genotyped as absent (grey area). In all cases, there is a sizeable overlap between data base variants discovered by DELLY and variants typed to be present by DIGTYPER (darker purple area). Variants in this set are most likely truly present in this individual. Combined these two areas of putatively correct results represent large fractions of the total variants and indicate 91.1% to 98.3% agreement. As is not uncommon for Venn diagrams of variant prediction methods, there is also a sizeable symmetric difference of variants typed to be present by DELLY or DIGTYPER but not both. The true status of these variants remains unknown to us, but one might hypothesize that not all these variants are false positives and that the two methods therefore complement each other.

## 4 Conclusion

In this paper, we have presented a new method to genotype tandem duplications and inversions. The issue in this is that the short read data that provides evidence of the genotype is affected by uncertainties, which can decisively hamper the task. Here, we have addressed this by a sound statistical framework that aims to determine the correct genotype as the most likely one given the short read data. It is common to maximum likelihood estimation procedures that naive approaches have exponential runtime when taking data uncertainties into account. One important achievement of ours has been to provide a computation scheme that allows to determine the genotype in runtime linear in the supporting short read data. As results, we have demonstrated that our method achieves significant improvements over DELLY, to date the only method that allows to genotype tandem duplications and inversions, in various aspects.

Still, there is room for improvements. For example, inversions, duplications, and deletions often come in combination, which we have not addressed here. Since our approach is flexible in terms of combining variants, we will be able to address also this case in the future. We consider it worthwhile to further invest in re-aligning reads so as to achieve refined alignment probabilities and even more accurate read alignments. Extending our read mapper LASER to also detect split alignments for duplications could potentially bring an improvement. Last but not least, marrying our duplication genotyping approach to coverage-based techniques is a promising future endeavor, for instance by using coverage signals to obtain priors.

